# Spatial distribution and habitat suitability of tsetse (*Glossina spp*.) in Côte d’Ivoire: An ensemble modeling approach to support targeted disease control

**DOI:** 10.64898/2026.08.21.746192

**Authors:** Antoine M.G. Barreaux, Harriet Wangu, Bamoro Coulibaly, Djakaridja Berté, Donatien K. Coulibaly, Elfatih M. Abdel-Rahman, Raphael Mongare, Pacôme Adingra, Vessaly Kalo, Stella Gachoki, Alain Boulangé, Geoffrey Gimonneau, Sophie Thevenon, Giuliano Cecchi, Philippe Solano, Dramane Kaba

**Author notes:** corresponding author: Antoine Barreaux. Equal contribution (Antoine M.G. Barreaux, Harriet Wangu).

## Abstract

**Background:** Tsetse are vectors of trypanosomes responsible for African animal trypanosomosis (AAT) and human African trypanosomiasis (HAT). While Côte d’Ivoire has successfully eliminated HAT as a public health problem and approaches elimination of transmission, AAT remains a major obstacle to agriculture and livestock production. Understanding the spatial distribution of tsetse is essential for prioritizing and sustaining disease control and elimination efforts.

**Methodology/Principal Findings:** Using 1,702 occurrence records from the national tsetse atlas we modeled the habitat suitability of the nine tsetse species present in Côte d’Ivoire. We identified suitable habitats in unsampled areas and quantified environmental constraints on tsetse distribution. Resampling the data to a 1km x 1km grid produced spatially explicit outputs at a resolution more relevant for operational planning. An ensemble modeling approach was employed integrating four algorithms—Random Forest, XGBoost, Maximum Entropy (MaxEnt), and Generalized Additive Models (GAM)— with satellite-derived environmental and anthropogenic predictors—which achieved high predictive accuracy, area under the curve and True Skill Statistics 0.80 and 0.83, respectively. Distance to waterbodies, soil moisture, distance to protected areas, maximum land surface temperature, and sheep density were key drivers of habitat suitability. Importantly, the models identified suitable habitats in 11 administrative regions not covered by the atlas, providing an improved national tsetse risk profile.

**Conclusions/Significance:** These results provide a detailed assessment of the ecological suitability of tsetse across Côte d’Ivoire and their persistence in agroecological mosaics with high human and livestock densities. We offer a high-resolution blueprint for vector and disease control, particularly in areas where field data are currently lacking. We provide a robust framework for evidence-based decision-making within the Progressive Control Pathway (PCP) for AAT by enabling the identification of priority areas and resource allocation optimization to improve livestock productivity through more effective AAT control and reduce the risk of resurgence of HAT.

**Author Summary:** Tsetse flies are a major threat to health across sub-Saharan Africa, transmitting parasites that cause sleeping sickness in humans and nagana in livestock. While Côte d’Ivoire has significantly reduced human cases, the impact on livestock still hinders economic growth. To protect both people and animals, health authorities need accurate maps showing where different tsetse species live. We analyzed over 1,700 records of tsetse from across Côte d’Ivoire, combining them with satellite-based data on temperature, vegetation, and water. Using machine-learning models, we created detailed maps predicting habitat suitability for nine tsetse species. Our findings reveal that different species confine to specific environments, with some living near rivers and others moving into areas with high human and livestock activity. These maps offer a high-resolution layer to inform HAT and AAT disease surveillance and control. By knowing where flies thrive, resources can be precisely allocated to high-risk areas. This study can help protect livestock, boost animal productivity, and ensures that human sleeping sickness does not return to regions where it was previously eliminated.

## Introduction

Tsetse fly (*Glossina spp.*) is a vector widely distributed across sub-Saharan Africa that transmits trypanosome parasites causing African animal trypanosomosis (AAT) and Human African trypanosomiasis (HAT). Due to its impact on agriculture and livestock production, AAT remains one of the main obstacles to socioeconomic development in affected rural areas across Africa (1,2). Approximately 100 million heads of cattle are estimated to be at risk of trypanosome infections (3), resulting in reduction in milk production by 10-40%, declines in cattle populations by 10-50% and a decrease in agricultural production by up to 2-10%. Collectively, the direct and indirect economic losses associated with AAT are estimated at US$5 billion annually (3). In contrast, the burden of HAT, a debilitating and deadly neglected tropical disease if left untreated, has declined substantially owing to highly effective control programs implemented over the recent decades, leading to official elimination of HAT as a public health problem in several endemic settings (4). However, sustained vigilance remains essential, as achieving and maintaining disease elimination is inherently challenging and requires continued surveillance to prevent re-emergence (4–7). Hence, the identification of tsetse habitat suitability, i.e. the type of natural environment in which the species can live or find food, shelter, protection and mates for reproduction (9), and their geographical distribution is essential for the targeting disease surveillance, control and potential elimination of both the animal and human diseases and their vectors (10–14).

The *Glossina* genus is taxonomically and ecologically divided into three groups based on morphological characteristics (specifically genitalia structure), habitat preferences, host associations, and geographical distribution: savannah (*morsitans* group), riverine (*palpalis* group), and forest species (*fusca* group) (15,16). This differentiation reflects the narrow dependence of tsetse on specific environmental and climatic variables. Both the male and female tsetse are obligate blood feeders, limiting their dependency on water for survival and enabling them to be present all year long. Tsetse fly employs a unique reproductive strategy known as adenotrophic viviparity—where larvae are nourished by specialized milk glands and pupate shortly after deposition hour (17–19). However, their survival remains strictly dependent on specific microclimates and host availability; hence, the spatial distribution of hosts, alongside water and vegetation cover, represents a critical ecological driver for predicting tsetse fly habitat suitability

Species distribution models (SDMs), also referred to as ecological niche models or habitat suitability models, assess the relationship between the species occurrence records and the environmental conditions associated with their geographic distribution. Increasingly, machine learning-based SDMs have been applied in numerous studies to assess vector habitat suitability using satellite-derived environmental predictors across spatial and temporal scales. Several studies modelled the suitability of tsetse fly (*Glossina spp.*) using SDMs (20–22) have demonstrated that these approaches are valuable not only for mapping the risk of HAT or AAT, but also for optimizing vector control operations (23). In particular, SDMs provide critical decision support tools for identifying priority intervention areas and guiding vector management strategies throughout the various stages of the progressive control pathway (PCP) for AAT (12,13,22). Moreover, extensive surveillance efforts have generated substantial occurrence datasets through continental and national tsetse atlases in several endemic regions (24–29), In such contexts, SDMs represent an essential next analytical step, enabling the extrapolation of observed occurrence patterns to identify potentially suitable habitats in non-surveyed or under-sampled areas that may require targeted monitoring and intervention. However, SDMs are inherently constrained by the assumption that observed presence records adequately represent a species’ current habitat preferences. In reality, these observations may capture only part of the species’ ecological niche and may not fully reflect its complete environmental tolerance or preferred ecotype. As a result, modeled distributions generally represent the realized niche, the portion of the niche occupied under current climatic, ecological, and biotic conditions, rather than the species’ full fundamental niche or potential geographic range. Furthermore, these realized distributions may be substantially modified by anthropogenic influences, including land-use change, habitat fragmentation, and vector control interventions, which can either suppress or preserve local populations.

In Côte d’Ivoire, HAT caused by *Trypanosoma brucei gambiense* has been eliminated as a public health problem (4) and is approaching elimination of transmission according to WHO criteria (30). Vector control has played a pivotal role in the achievement of these milestones (31), and a better understanding of tsetse fly distribution remains important for preventing disease resurgence and mitigating potential risks associated with the hypothesized animal reservoir (8,32). In northern Côte d’Ivoire, however, tsetse and AAT continues to pose a major constraint to livestock health and agricultural productivity, national meat self-sufficiency and expansion of key cash crops such as cotton, which depend heavily on for animal draught power (33). Recent molecular surveillance on cattle blood samples (33) using Polymerase Chain Reaction (PCR) has revealed persistently high infection levels, with an overall prevalence of approximately 12.3%, substantially exceeding estimates obtained by conventional microscopy [approximately 5%]. The departments of Korhogo, Boundiali, and Ferkessédougou remain the most severely affected areas. Given the continued persistence of the disease despite existing control measures, future intervention strategies should adopt a One Health approach that integrates high-resolution habitat suitability modeling, community-based vector control and improved livestock management to enhance disease control, strengthen livestock productivity, and safeguard national meat self-sufficiency and animal traction systems.

In Côte d’Ivoire all three ecological groups of tsetse fly are present: savannah (*G. longipalpis* and *G. morsitans submorsitans*), riverine (*G. tachinoides, G*. *p. palpalis, G. palpalis gambiensis*, and *G*. *pallicera*) and forest species (*G*. *medicorum*, *G. fusca fusca,* and *G*. *n. nigrofusca*), occupying diverse ecological zones ranging from the dense humid forests in the south to savannah and transitional habitats in the north (34). However, recent environmental changes, driven by deforestation, the transition to cocoa and coffee plantations, agricultural expansions, settlement patterns that involve livestock keeping (including pigs, goats, sheep, and cattle), and climate variability, have significantly altered tsetse fly habitats and host availability. The recent development of a national atlas (24) has provided a comprehensive baseline of tsetse fly across Côte d’Ivoire, creating a valuable foundation for ecological niche modelling to predict the potential distribution of vector species and identify high-priority areas where entomological surveillance remains limited or absent.

In this study, we aimed to predict the current (2018–2023) habitat suitability of nine tsetse fly species (*Glossina spp*.) in Côte d’Ivoire using an ensemble ecological niche modelling approach. We first conducted a comprehensive literature review and consulted expert knowledge to identify the key and relevant environmental drivers influencing tsetse fly distribution. These variables were subsequently used to develop an ensemble of multiple machine learning algorithms that integrated tsetse fly occurrence records with environmental predictors to model the ecological niches and predict the potential distribution of each species across the country. The resulting habitat suitability maps provide a spatially explicit evidence base to support decision-making, optimize resource allocation, and prioritize surveillance, vector control, and field interventions. These outcomes primarily contribute to Stage 1 of the Progressive Control Pathway (PCP) for AAT, which focuses on establishing information systems, assessing disease risk and impact, and identifying priority intervention areas. They also support Stage 2 by informing targeted and sustainable strategies to reduce the burden of AAT (35).

## Methods

We developed SDMs for the nine tsetse fly species present in Côte d’Ivoire using their occurrence records from the national tsetse fly atlas (24) combined with explanatory environmental variables derived from open-access datasets and satellite-based remote sensing variables (36–40). Fig. 1 below shows the integration and modelling process of the whole process.

**Fig. 1.**
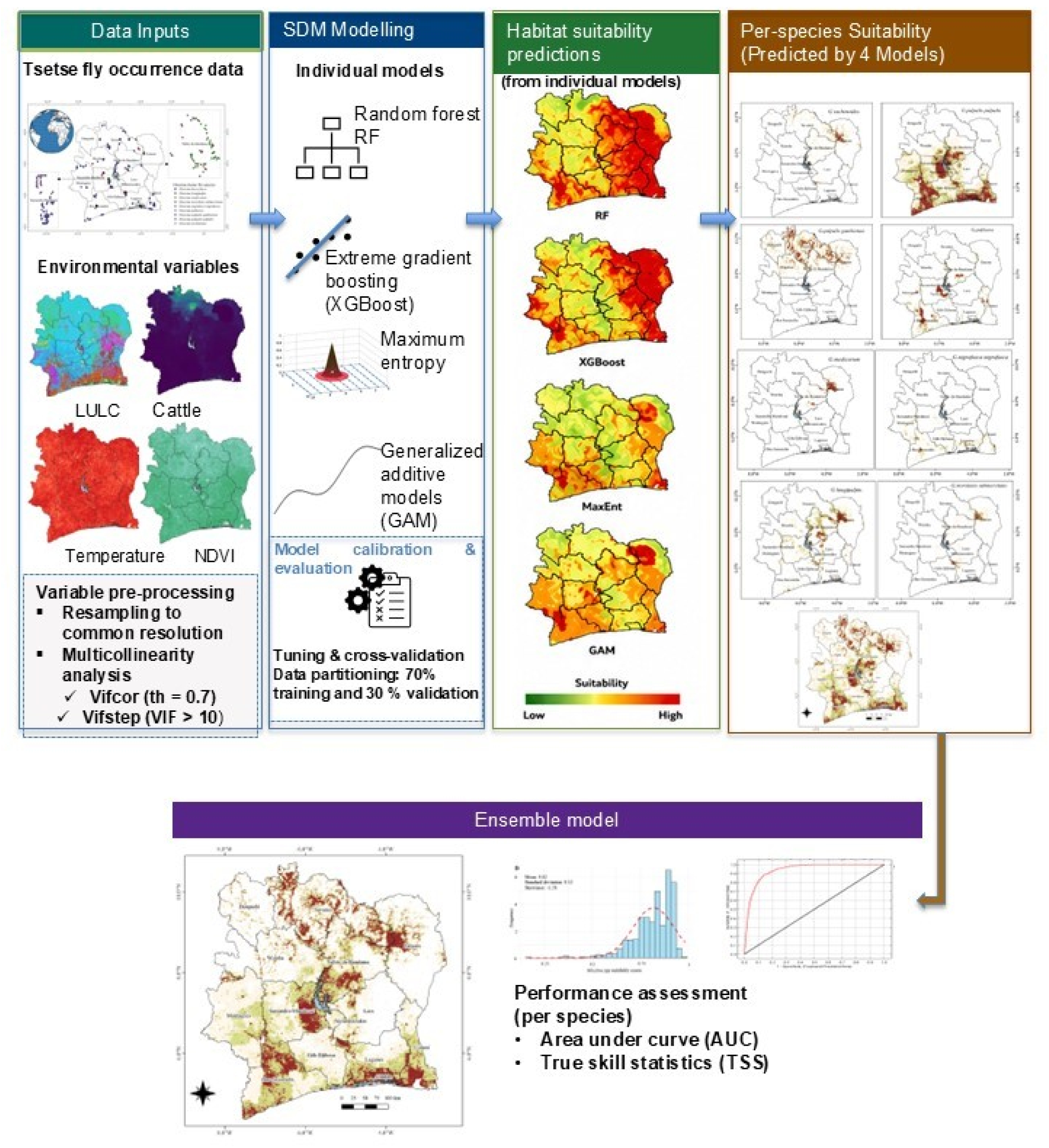
The general methodological framework employed in this study to predict the habitat suitability of tsetse fly (*Glossina spp.*) in Côte d’Ivoire. The four species distribution models used are random forest (RF), extreme gradient boosting (XGBoost), maximum entropy (MaxEnt) and generalized additive model (GAM). The ensemble was generated using an ensemble machine learning modelling approach based on true skill statistic [TSS]-weighted mean. The environmental variables abbreviated as LULC and NDVI refer to land use/land cover and normalized difference vegetation index, respectively.

### Study area

Côte d’Ivoire’s varied habitats (Fig. 2) range from tropical rainforests in the southern and savannas in the middle and northern areas to coastal lagoons and mangroves in the southern part of the 322,463 km² area (41). There are various humid, sub-humid, and dry zones in the mostly tropical climate, which affects land usage, biodiversity, and the habitats of disease vectors.

**Fig. 2.**
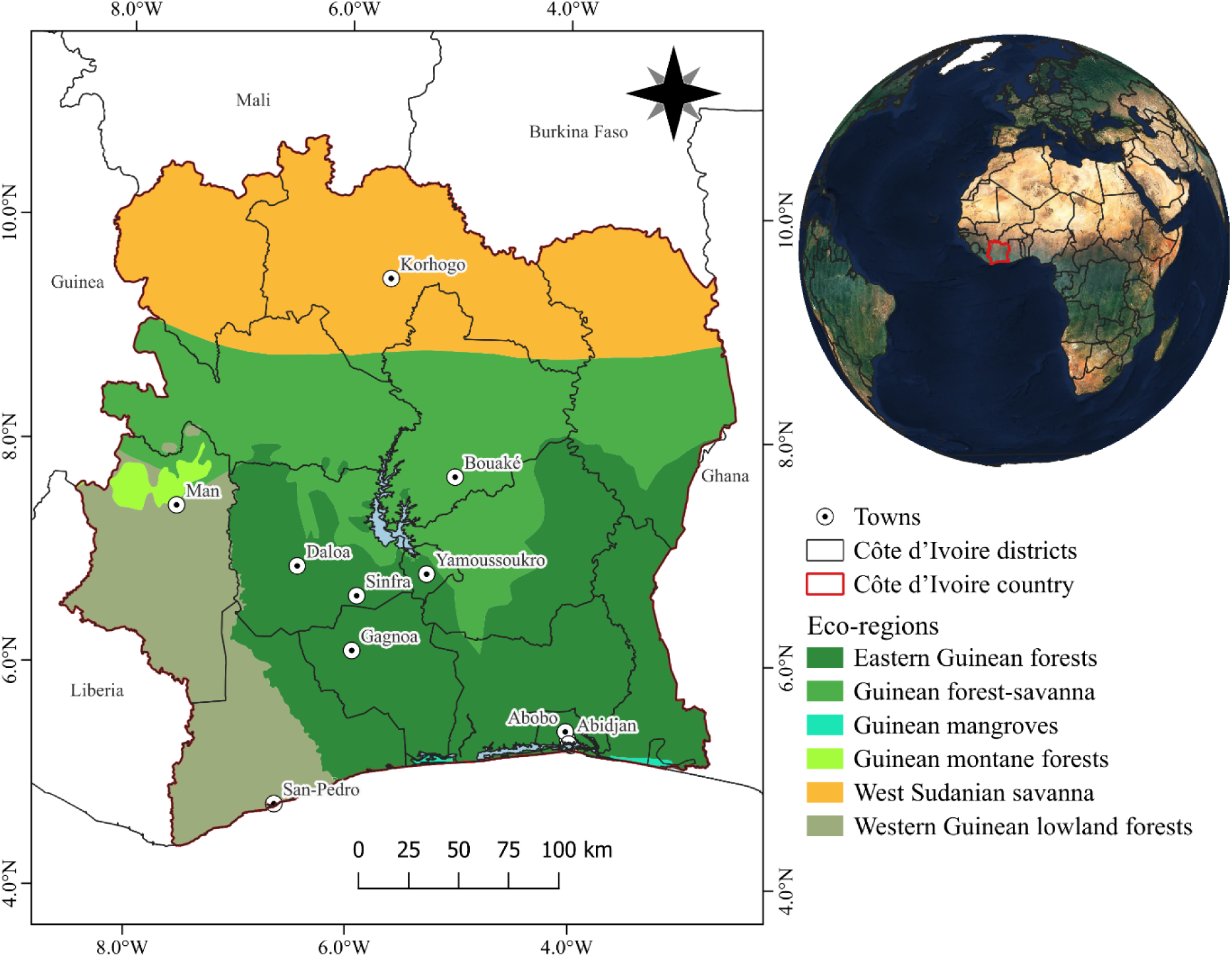
Study area showing the location of Côte d’Ivoire in West Africa, its administrative districts, major towns, and the distribution of the six ecoregions: Eastern Guinean forests, Guinean forest–savanna mosaic, Guinean mangroves, Guinean montane forests, West Sudanian savanna, and Western Guinean lowland forests.

Côte d’Ivoire has mean annual temperatures ranging from 25 to 27 °C across the country. Annual rainfall ranges from 1000 to 1600mm with higher amounts in the north and south and lower amounts in the central part of the country (42). In this climate zone, four seasons are generally distinguished: The large rainy season from April to the end of July, the small dry season from August to September, the small rainy season from October to November, and the large dry season from December to the end of March. In the central part of country, the annual temperatures are between 20 and 40°C. The seasons are roughly the same as on the coast, but there is less rainfall. In the northern part the annual temperatures are between 10 and 41°C (43).

### Tsetse occurrence data and sampling techniques

The tsetse fly occurrence dataset used in this study comprised field reference points obtained from the Côte d’Ivoire tsetse fly national atlas (24) focusing on nine species of tsetse flies (*Glossina* spp.), representing three ecological groups: savannah species (*G. longipalpis* and *G*. *morsitans submorsitans*), riverine species (*G. tachinoides*, *G*. *p. palpalis*, *G. p. gambiensis*, and *G. pallicera*) and forest species (*G. medicorum*, *G. fusca fusca,* and *G. n.nigrofusca*) distributed across Côte d’Ivoire. All tsetse fly survey records collected by the Institut Pierre Richet (IPR) and collaborators between January 2005 and December 2024 were compiled, georeferenced and harmonized into a national occurrence database. These records originated from 20 of the country’s 31 administrative regions. For the SDM, we focused on the most recent five-year period (2018–2023), which provided both a high density of occurrence observations and access to high-quality Earth observation datasets for environmental characterization. In addition, *G. fusca fusca* was excluded from species-specific habitat suitability modelling because of the limited number of occurrence points and their restricted spatial distribution (Fig. 3). In total, 3,164 trapping sites were recorded, representing 5,696 individual trapping events and a cumulative sampling effort of 13,181 trap days. Following data cleaning, including the removal of records with missing values and duplicate observations, the final dataset compromised 1,702 unique occurrence records used for model development (see S1 Dataset for more information). Fig. 3 shows the distribution of the tsetse fly occurrence points based on ecological groups used in the SDMs.

**Fig. 3.**
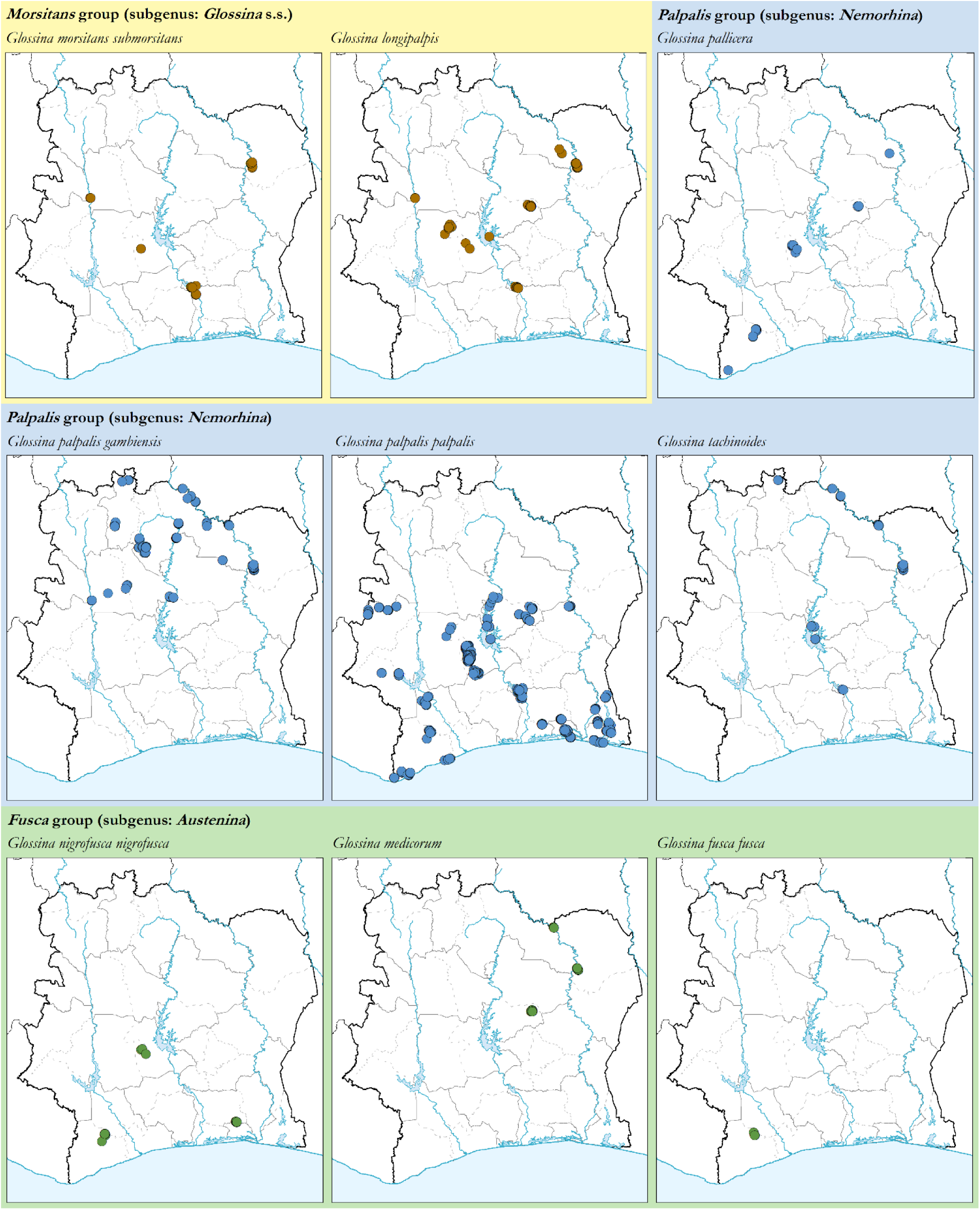
Tsetse fly (*Glossina* spp.) occurrence records representing three ecological groups collated across Côte d’Ivoire from 2018 – 2023.

### Dependent explanatory variables

After a comprehensive literature review and expert knowledge consultation, several environmental variables were selected based on their relevance to tsetse ecology and behavior, as summarized in Table 1. The dependent variables covered the period between 2018 and 2023, consistent with the tsetse fly presence records. The variables include climatic, topographic, vegetation density, land use/ land cover (LULC), distance to protected areas and demographic drivers, that have previously been used in determining tsetse fly habitat suitability (20). The LULC layer (37) included several classes (labels) that were described in (S1 appendix, Table S1). These dependent variables were obtained from different sources with varying spatial resolutions. We use the Google Earth Engine platform (44) to obtain and harmonize the resolution of the variables to (1 km x 1 km) grid cell. Regarding the distance to protected areas, we used the proximity function in QGIS software (QGIS Development Team, 2024) to calculate the distance of the tsetse fly occurrence points from the protected areas and water bodies (see S1 Folder_Environmental covariates data to explore these layers).

**Table 1.** The dependent variables, along with their original spatial resolution, units, sources, and rationale for inclusion in assessing the distribution of tsetse fly in Côte d’Ivoire.

| Variable | Spatial resolution | Unit | Source | Rationale |
| --- | --- | --- | --- | --- |
| Precipitation | 4.6383 km<sup>2</sup> | Millimeters | https://www.climatologylab.org/terraclimate.html (45) | Precipitation influences humidity and the availability of water resources, critical for tsetse fly and host survival (46). |
| Land surface temperature | 1 km<sup>2</sup> | Kelvin | https://lpdaac.usgs.gov/products/mod11a1v061/ (47) | Temperature affects tsetse fly metabolism, development, fecundity and survival rates, where tsetse fly survives below 33°C and above 16°C (48–50) and may also be key to vectorial capacity and hence disease transmission (14) |
| Normalized difference vegetation index (NDVI) | 500 m<sup>2</sup> | Meters | https://lpdaac.usgs.gov/products/mod13a1v061/ (51) | NDVI indicates vegetation health and density, which influence tsetse fly habitat suitability and resting sites (48). |
| Elevation and aspect | 30 m | Meters and degrees | https://lpdaac.usgs.gov/products/srtmgl1v003/ (52) | Elevation and slope orientation affect microclimate, temperature, and moisture availability for tsetse fly. |
| Land use/ land cover (LULC) | 10 m | Meters | https://africageoportal.maps.arcgis.com/apps/webappviewer/index.html?id=88c2493e722546c09c2a0a8b394c4454 (37) | LULC determines tsetse fly habitat type, human impact, and availability of host species and breeding sites (53). |
| Waterbodies | 10 m | Meters | https://africageoportal.maps.arcgis.com/apps/webappviewer/index.html?id=88c2493e722546c09c2a0a8b394c4454 (37) | Distance to water bodies acts as a proxy for microclimates that are suitable for the tsetse and hosts. Such conditions include high humidity and moderate temperatures (54). |
| World Protected areas | *NU | Kilometer | https://www.protectedplanet.net/en/thematic-areas/wdpa?tab=WDPA (40) | Tsetse occurrence is often higher near protected areas due to the presence of abundant hosts and favorable environmental conditions such as shaded habitats. As distance from these areas increases, tsetse populations generally decline. However, this relationship may vary depending on species ecology and local environmental conditions (48). |
| Soil moisture | 4.6383<br>km<sup>2</sup> | Millimeters | https://www.climatologylab.org/terraclimate.html (45) | Soil moisture affects tsetse fly larval and pupal development in soil. Highly suitable moisture content holding provides suitable micro-climate for tsetse fly (15). |
| Livestock density (cattle, sheep, goats, pigs) | km<sup>2</sup> | Number of livestock per square kilometer | https://www.fao.org/livestock-systems/global-distributions/en/ (39) | Livestock serves as a key blood meal host, influencing tsetse feeding and survival (55). |
| Human population | ~Approximately 1<br>km<sup>2</sup> | Population density | https://www.earthdata.nasa.gov/data/projects/gpw (38) | Human population density affects habitat disturbance, availability of hosts, and potential disease spread risk (56). |

\*NU – no spatial resolution

### Exploratory data analysis

Multicollinearity among the environmental predictors was assesses using the “vifcor” and “vifstep” functions implemented in the usdm R software (57,58). First, the “vifcor” function was used to identify highly correlated predictors pairs based on Pearsons correlation coefficients Following Dormann (59) an absolute correlation threshold of (r ≥ 0.7) was implemented. For each correlated pair, the variables with the higher variance inflation factor (VIF) were removed iteratively until no remaining predictors pairs exceeded the specified correlation threshold. Next the “vifstep” function was applied to assess any residual multicollinearity among the remaining predictors. Variables with VIF values greater than 10 were considered highly collinear and were sequentially removed until all retained variables had VIF values < 10. A total of 24 out of 66 initial environmental predictor variables were selected and retained for predicting the habitat suitability of tsetse fly based on the VIF and correlation diagnostics (S1 appendix, Table S2). The retained variables exhibited substantially reduced collinearity, with linear correlation coefficients ranging from a minimum of 0.00027 to a maximum of 0.69105 (S1 appendix, Table S2). The Spearman correlation matrix showed the pairwise relationships among the retained predictors (S1 appendix, Fig. S1). In addition, hierarchical cluster analysis grouped the 24 retained variables into five major clusters according to the similarity of their spatial information (Fig. 4). The five clusters indicated distinct but related environmental dimensions within the predictor set, with groups of variables exhibiting similar spatial characteristics across the study area.

**Fig. 4.**
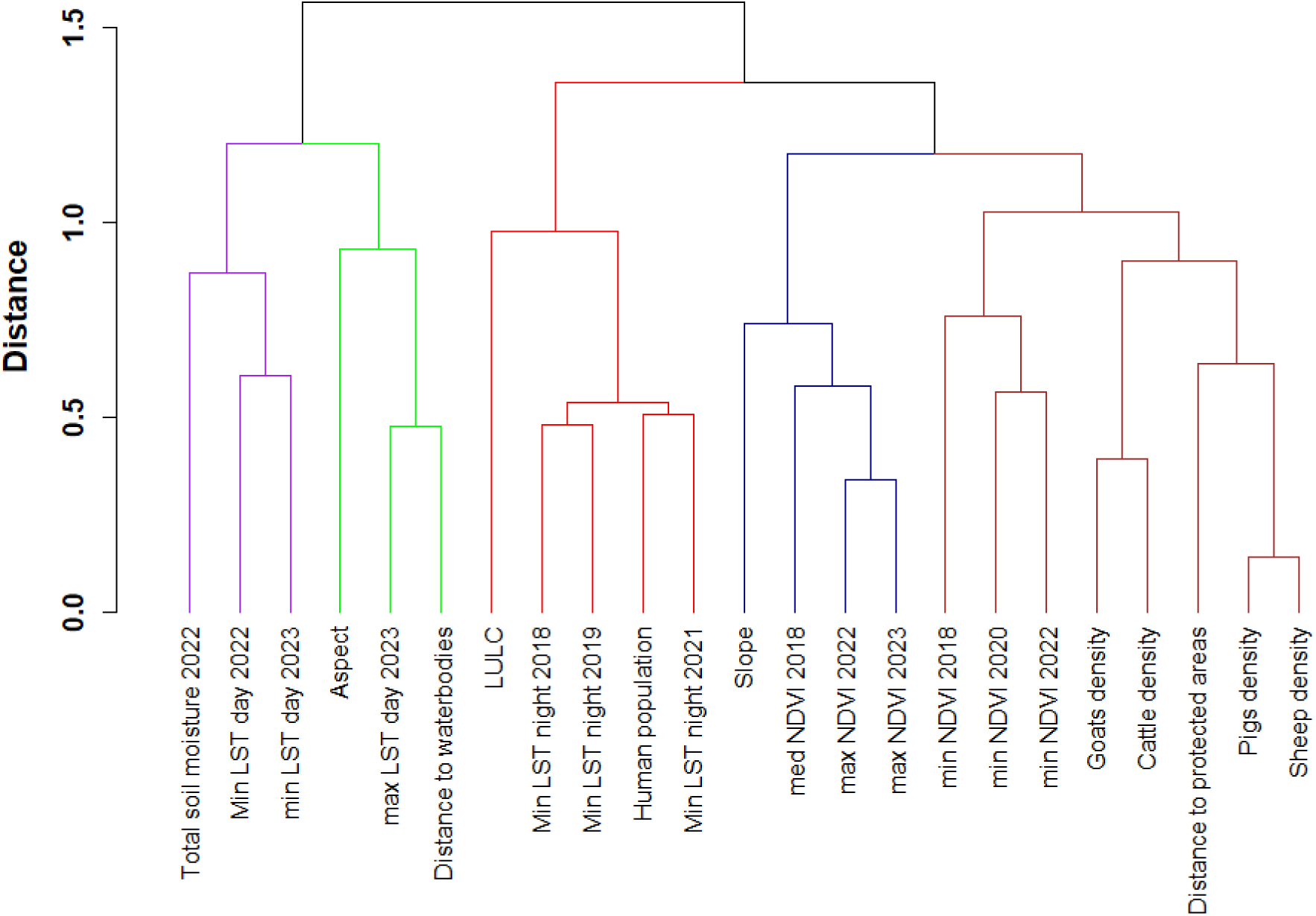
Hierarchical clustering of the environmental variables retained after VIF analysis. The dendrogram illustrates the degree of similarity (distance) among variables. Variables grouped within the same-colored cluster at linkage distances of 0–0.5 exhibit strong correlations, those clustering between 0.5–1.0 show moderate correlations, whereas variables joined at linkage distances of 1.0–1.5 display relatively weak correlations.

### Species distribution modeling [SDM] and ensemble approach

Species distribution modelling (SDMs) are numerical tools that relate observations of species occurrence or abundance to environmental conditions for predicting the spatial distribution of the species. They are used to gain ecological and evolutionary insights about the species and to predict their distributions across landscapes (60). In this study, we used four different machine learning SDM algorithms viz., random forest (RF) (61), extreme gradient boosting (XGBOOST) (62), maximum entropy (MaxEnt) (63) and generalized additive model (GAM) (64) in the biomod2 package (65). Each of these algorithms has unique strengths, weaknesses, and underlying assumptions when applied to SDM (S1 appendix, Table S3). The MaxEnt model is a machine learning method that uses presence-only data to estimate the species habitat suitability by finding the most uniform distribution constrained by environmental conditions at known species locations (63). Random forest (RF) is an ensemble method that combines multiple decision trees to handle complex ecological data, providing robust classification and regression outputs along with measure of variable importance (66). Whereas GAM is a flexible data-driven approach that fits smooth, non-linear relationships between species presence and environmental variables without assuming predefined parametric forms (67). XGBoost is a powerful gradient boosting algorithm that sequentially builds decision trees, each correcting the errors of the previous one to optimize predictive accuracy (68).

The potential distribution of multiple *Glossina* species was predicted using an ensemble SDM framework implemented in the R package biomod2 (65). The modeling workflow was automated through a custom function designed to process each species individually, incorporating occurrence data, environmental predictors, and multiple algorithms to generate robust distribution predictions. Occurrence records for each *Glossina* species were filtered to remove any records with missing geographic coordinates or incomplete data. A 1 km grid was generated to spatially separate presence and absence points and reduce spatial autocorrelation. However, only seven absence points were retained in unique grid cells independent of presence locations (out of the 47 total absence points), indicating a highly limited and spatially biased absence dataset. Consequently, a presence-only modelling approach was adopted to ensure robust and reliable habitat suitability predictions with 1,655 remaining occurrences. For each species, presence points were extracted, and environmental variables were sampled at these locations from a raster stack (S1 Folder_Environmental covariates data).

To complement presence-only data, pseudo-absence points were generated randomly within the study area using a random sampling strategy. For each species, 1,000 pseudo-absence points were created, ensuring they did not overlap with known presence locations. This approach helps balance the presence-absence dataset and improves model reliability. Species data were formatted for biomod2 using the BIOMOD_FormatingData function, specifying the response variable (presence = 1, pseudo-absence = 0), explanatory environmental variables, and geographic coordinates. Five repetitions of pseudo-absence generation were performed to account for uncertainty in absence data.

Model calibration was performed using a random cross-validation strategy with 70% of the data used for training and 30% for testing, repeated five times to assess model stability. The optimization strategy “bigboss” was applied to tune model parameters automatically. Table 2 shows the summary of the parameters, this strategy applies iterative hyperparameter optimization to systematically explore a wide range of model settings and identify parameter combinations that maximize predictive performance, considering key aspects of SDM (65).

**Table 2.**
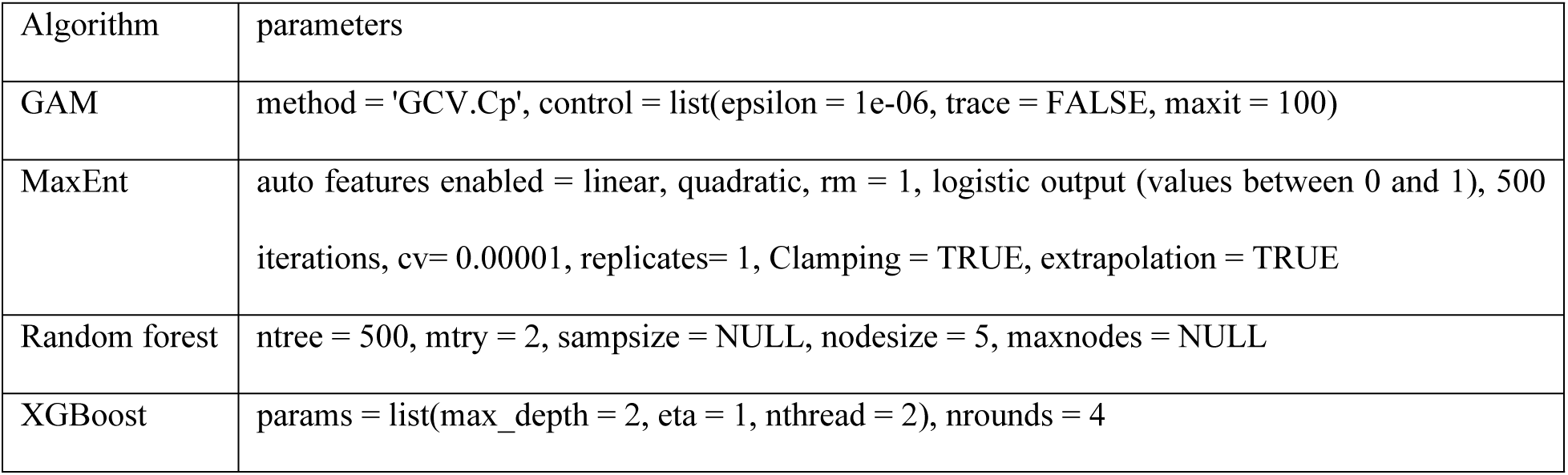
The bigboss tuned parameters used in each of the four machine learning species distribution models.

| Algorithm | parameters |
| --- | --- |
| GAM | method = 'GCV.Cp', control = list(epsilon = 1e-06, trace = FALSE, maxit = 100) |
| MaxEnt | auto features enabled = linear, quadratic, rm = 1, logistic output (values between 0 and 1), 500<br>iterations, cv= 0.00001, replicates= 1, Clamping = TRUE, extrapolation = TRUE |
| Random forest | ntree = 500, mtry = 2, sampsize = NULL, nodesize = 5, maxnodes = NULL |
| XGBoost | params = list(max_depth = 2, eta = 1, nthread = 2), nrounds = 4 |

The variable importance for predicting tsetse fly distribution was assessed by permuting each environmental variable and measuring the influence on model predictions, providing insights into the relative contribution of variables across algorithms. Only models meeting certain performance thresholds (true skill statistic: TSS and area under curve: AUC > 0.6) were retained for the ensemble modeling experiment (65,69,70).

Multiple ensemble methods are available, including mean, weighted mean, and median, as consensus approaches, to combine individual model predictions and reduce uncertainty. In this study, a weighted (by TSS) mean ensemble approach was applied to combine individual model predictions and to generate habitat suitability maps for *Glossina spp.* under current environmental conditions. All analyses were conducted in R (version 4.5.1) to ensure reproducibility (57,58). The modeling procedure was applied iteratively to all target *Glossina* species, enabling comparative assessment of distribution patterns and environmental drivers. The code of the whole approach in R can be found in “S3 Appendix_Species distribution modeling code”.

### Evaluation of model performance

The performance of the SDMs was evaluated using the AUC. The AUC indicates how well a model can discriminate between species suitable areas and areas that are not suitable, with a suitable value range from 0.5 to 1. We also used TSS, which corrects for dependence on species prevalence while assessing how the model differentiates between the species presence and absence records (69).

We further validated the performance of models using an independent species-specific test dataset for *G*. *p. palpalis* collected in 2024 and a dataset at genus level (*Glossina* spp.) collated between 2007-2017 (older data that the data used for model development) from the Côte d’Ivoire national atlas (24). All the test datasets were georeferenced occurrence points. The test data points were overlaid on the *G*. *p. palpalis* habitat suitability map to extract the habitat scores that were used to fit histogram and normal distribution curves. The mean, standard deviation and skewness values were then calculated to assess the model performance.

Committee averaging was used to summarize predictions from the selected models by first converting each model’s probabilistic output into a binary presence-absence prediction using the optimal threshold defined during model calibration (i.e., the threshold that maximizes the evaluation metric during the BIOMOD_Modeling step). Each model then “votes” on whether the species is present (1) or absent (0) at a given location, and the committee averaging score is calculated as the proportion of models voting for presence. This voting-based approach not only generates a consensus prediction but also provides an explicit measure of uncertainty: values close to 0 or 1 indicate strong agreement among models, whereas values around 0.5 reflect high disagreement.

## Results

### Collinearity test and variable selection

The correlation analysis between the environmental variables at the tsetse fly (*Glossina spp*.). occurrence [n =1,655] points enabled selection of key uncorrelated variables that were included to further assess the suitability for Tsetse fly (*Glossina spp*.) in Côte d’Ivoire. Hierarchical cluster analysis was used to visualize the similarity among the retained predictors, grouping them into five major clusters based on their correlation patterns (Fig. 4). The clustering indicates that variables within the same group share greater similarity than those in different groups, while all retained predictors satisfied the VIF threshold and were therefore included in the subsequent modelling analyses.

### Models’ accuracy, comparison, validation and uncertainty

The model accuracy results shown correspond to the mean weighted ensemble models for the individual species. RF and XGBoost consistently achieved higher AUC > 0.8 scores and TSS of > 0.7 across most tsetse fly species (Fig. 5), suggesting that they are more accurate in predicting tsetse fly habitat suitability. Based on the model performance metrics, only models that scored TSS > 0.6 were included in the ensemble modelling approach.

**Fig. 5.**
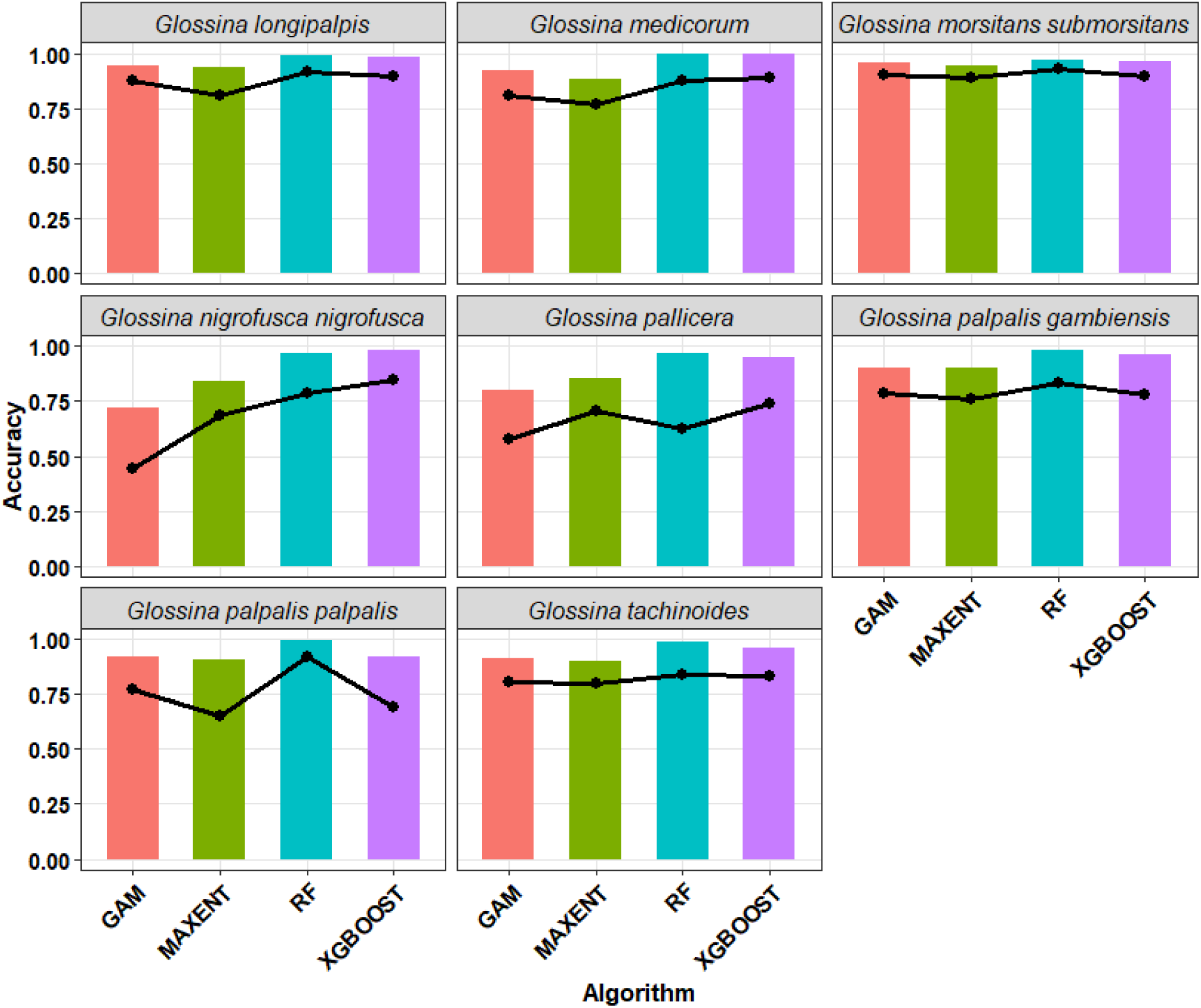
Validation performance of the four machine learning species distribution modelling algorithms (Generalized Additive Models (GAM), Maximum Entropy (MaxEnt), Random Forest (RF), and Extreme Gradient Boosting (XGBoost)) in predicting habitat suitability of nine tsetse fly (*Glossina spp*.) species in Côte d’Ivoire. AUC = Area Under Curve, TSS: True Skill Statistic. The AUC values, depicted by the height of the bars, indicate the models’ overall ability to correctly classify presences and absences of tsetse fly records, with higher scores representing better discrimination. The TSS values displayed as colored lines, measure the models’ accuracy with higher values reflecting improved predictive skill.

For *Glossina p. palpalis* (Fig. 6A), the suitability scores across modeled locations tend to cluster near the upper range (0.8-1), with mean of 0.83, and negative skewness (−0.88). This indicates that most validation points occurred in areas predicted to have a high habitat suitability. Similarly, the ensemble suitability scores for all *Glossina spp.* (Fig. 6B) had a mean value of 0.82 and showed stronger negative skewness (−1.58), suggesting that most areas were assigned relatively high suitability scores. Overall, these results suggest that most predicted habitats are suitable for *Glossina* flies.

**Fig. 6.**
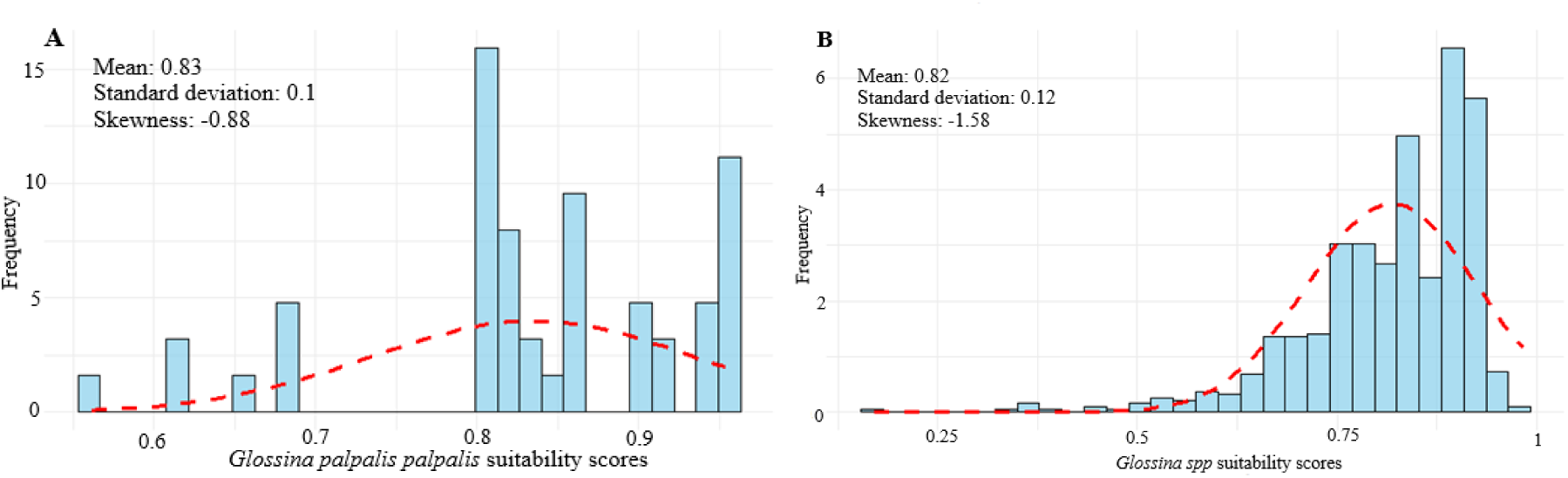
Histograms depicting the distribution of habitat suitability scores for *Glossina palpalis palpalis* (A) and for *Glossina* species (B). The fitted curve (red dashed line) highlights the overall distribution pattern of the *Glossina palpalis palpalis* or the *Glossina spp.* ensemble models.

Lower uncertainty values of the predicted habitat suitability for tsetse fly are observed in the southern Zanzan, Lacs, and parts of Woroba and Denguélé districts, where committee-averaging values approach 0 (Fig. 7). This pattern indicates strong model agreement that these areas are unsuitable for *Glossina* species, reflecting consistent predictions across all *Glossina* species models. In contrast, districts such as eastern Sassandra-Marahoué, southern Vallée du Bandama, parts of Savanes, Abidjan, Lagunes, and western Zanzan show high model agreement and therefore higher certainty in predicting tsetse fly habitat suitability.

**Fig. 7.**
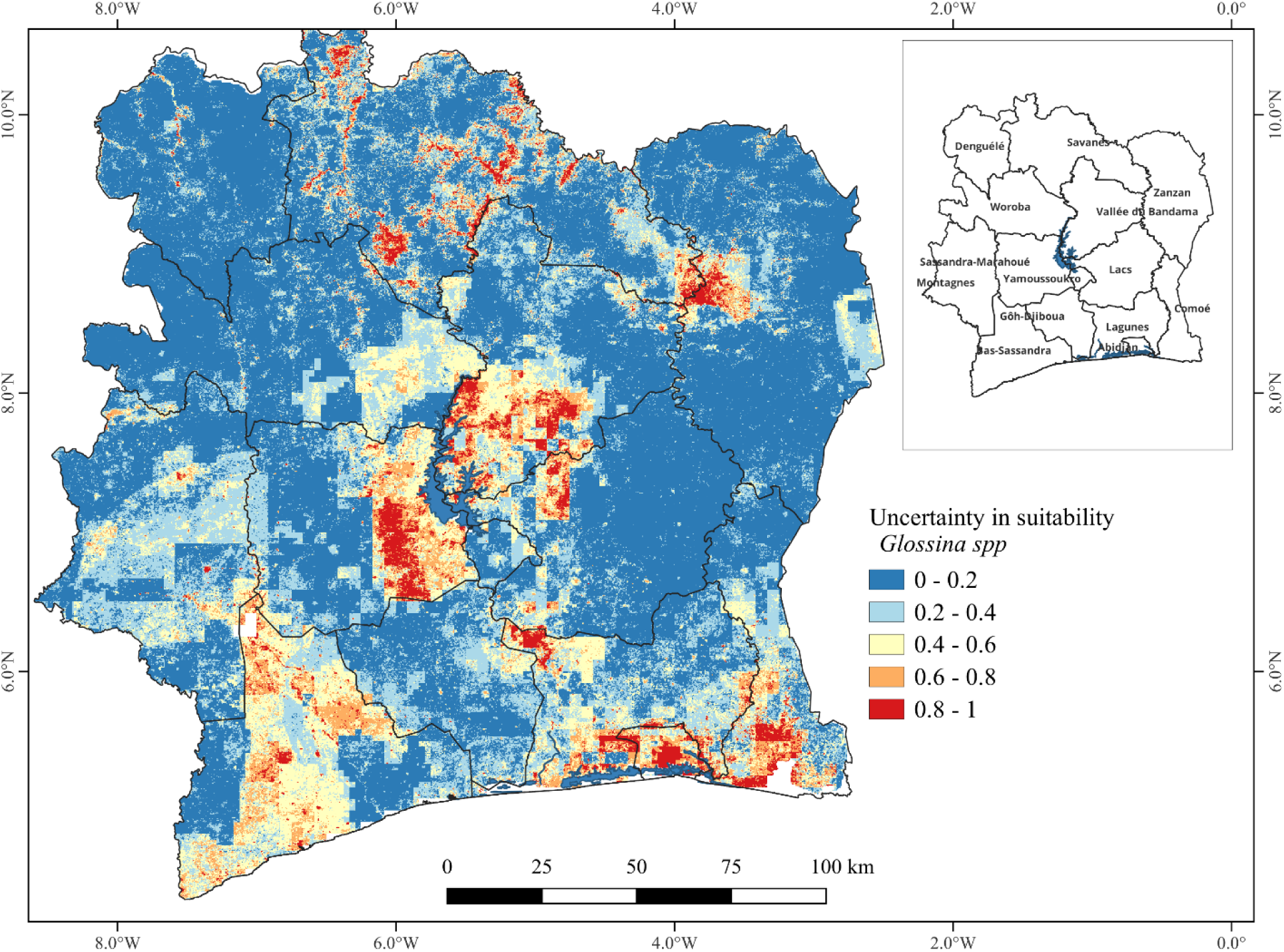
Model uncertainty in predicting tsetse fly (*Glossina spp*.) habitat suitability in Côte d’Ivoire using committee averaging (EMca). When the prediction is close to 0 (blue) or 1 (red), it means that all models agree to predict 0 and 1, respectively, and when the prediction is around 0.5, it means that half the models predict 1 and the other half predicted 0 indicating uncertainty in the predictions.

Moderate committee-averaging values (0.4-0.6) in the Bas-Sassandra district indicate disagreement among ensemble members, with approximately half of the model predictions (across nine *Glossina* species and multiple algorithms) favoring presence and half favoring absence. This pattern aligns with expectations given the diverse ecological preferences of the species included in the ensemble.

### Variable importance and response curves

Certain environmental and host-related variables such as distance to waterbodies, total soil moisture, distances to protected areas, maximum LST, and sheep and cattle densities (livestock densities overall) consistently showed high importance scores, especially within the GAM algorithm (Fig. 8). These variables likely play a pivotal role in defining suitable habitats for tsetse fly. Comparing across algorithms, it is evident that GAM often assigns pronounced importance to several predictors, while MaxEnt, RF, and XGBoost distribute their scores more evenly among the variables.

**Fig. 8.**
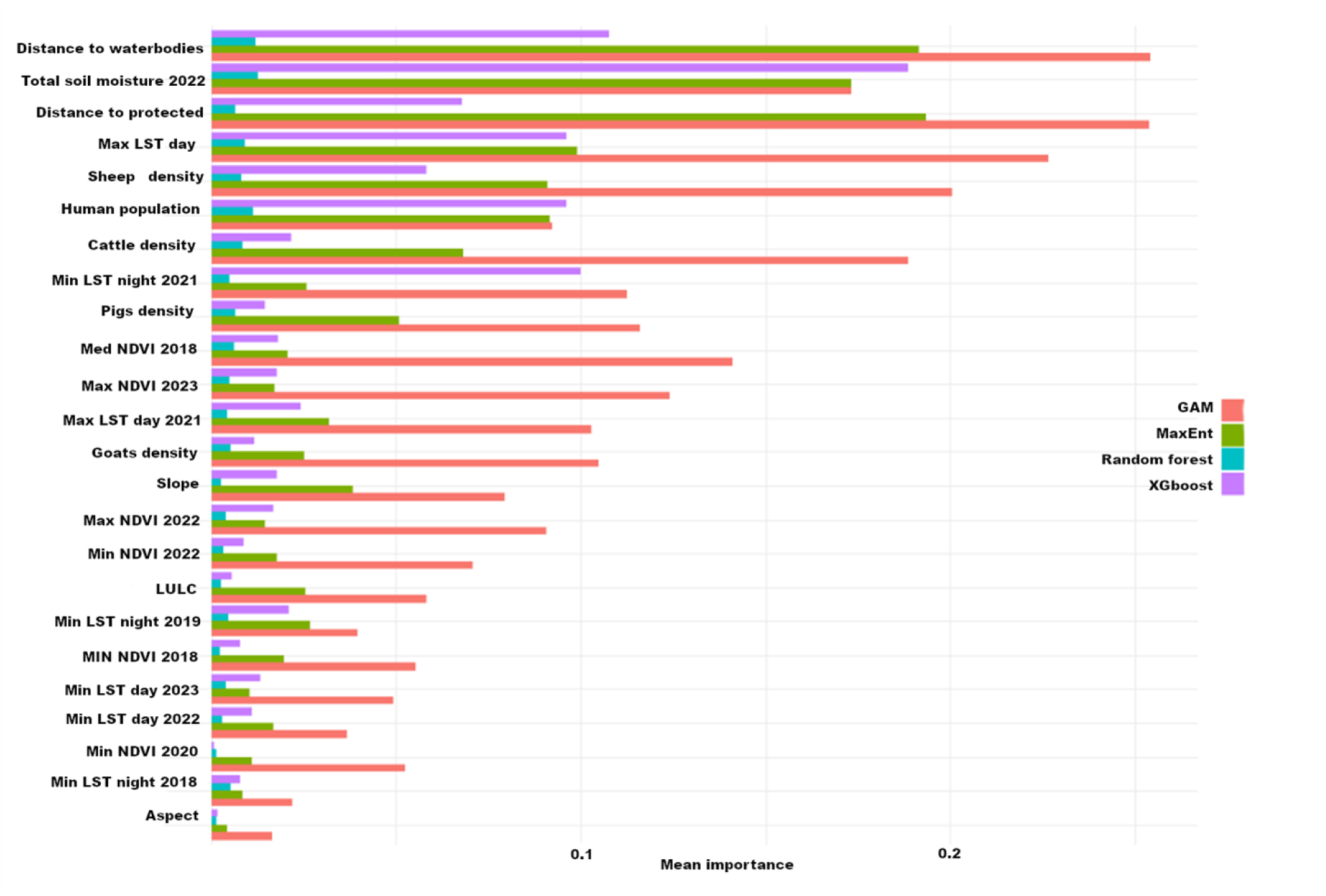
Mean importance scores of various predictor variables in predicting tsetse fly (*Glossina spp*.) distributions in Côte d’Ivoire based on weighted mean ensemble True Skill Statistics (TSS) model across different machine learning algorithms (Generalized Additive Models (GAM), Maximum Entropy (MaxEnt), Random Forest (RF), and Extreme Gradient Boosting (XGBoost)).

At the species level (S1 appendix, Fig S2-9), the variable importance patterns reinforce the genus-wide trends while revealing ecologically meaningful distinctions among tsetse groups. Vegetation greenness, particularly NDVI-based metrics, consistently emerges as a dominant predictor for several species such as *G. longipalpis*, and *G. n. nigrofusca*, emphasis the importance of humid, shaded habitats for these savannah and forest-associated taxa. Livestock density variables including cattle, goats, pigs, and sheep, also frequently attain high to moderate importance across species, especially in MaxEnt and GAM models, bringing out the role of host availability in shaping tsetse occurrence. Distance to waterbodies becomes particularly influential for riverine species (*G. p. palpalis*, *G. tachinoides*) and forest species, *G. n. nigrofusca,* aligning with their dependence on moist and high humidity forest environments, which places them near waterbodies and shaded wetlands. Seasonal LST metrics contribute moderately across species.

The response curves in Fig. 9 for *Glossina spp.* ensemble model showed that suitability increases with human population density (0-30,000 people km^−2^ and remains stable after that with a high suitability) and when the minimum values of LST go from 280 to 290 kelvin (7-17°C). Whereas habitat suitability decreases with an increase in livestock density. The LULC classes (water bodies, rivers, and waterways, wetlands/swampy areas and human settlements/infrastructure), higher NDVI indexes, increase in distance from protected areas and increase in distance from waterbodies have negative effect on *Glossina spp*. habitat suitability.

**Fig. 9.**
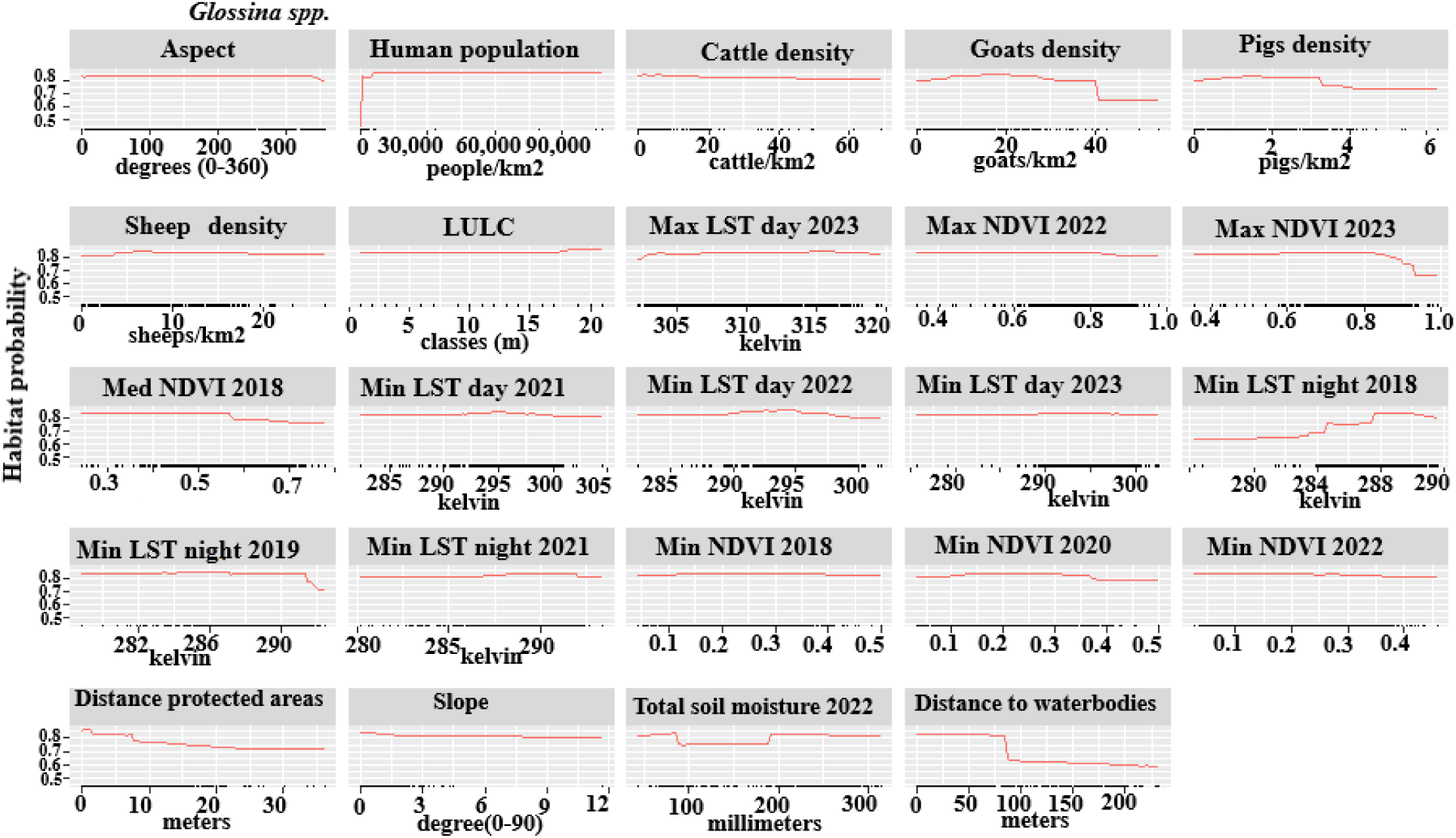
Response curves for the predictor variables in the final tsetse fly (*Glossina spp.*) habitat suitability model. The curves illustrate the relationship between each predictor variable and predicted habitat suitability.

The response curves (S2 appendix, Fig S1-8) for *G. longipalpis*, *G. n. nigrofusca*, and *G. pallicera* indicated that habitat suitability increases sharply with rising NDVI values, showing their dependence on dense, humid vegetation. Also, proximity to waterbodies produces strong positive responses for riverine taxa including *G. p. palpalis, G. tachinoides*, and *G. p. gambiensis*, whose suitability increases at very short distances from waterbodies. Livestock density variables [cattle, goats, sheep, pigs] also generate positive responses for several species, particularly the palpalis and morsitans groups, highlighting the significance of host presence in shaping habitat favorability. Human population density, in turn, elicits moderate to high suitability responses in species associated with changing landscapes due to human activities, especially *G. pallicera* and *G. p. gambiensis*, suggesting their adaptability to areas with intensive human–livestock interaction.

The LULC showed unique influence on the species, i.e. for *G. medicorum* is primarily associated with classes dense/open/gallery/secondary Forest, Forest plantation/reforestation/swamp forest, coffee/cocoa/rubber plantation, while suitability declines (below 0.5) in oil Palm/Coconut/Cashew/Fruit/Orchards/Agricultural land plantation, Tree savanna (wooded savanna), Shrubland/thickets, Herbaceous formations, waterbodies, and human settlements/infrastructure classes.

Climatic variables, such as seasonal minimum LST and daytime LST metrics, show optimum responses across species, suggesting the existence of optimal thermal envelopes rather than linear relationships. *G. medicorum*, *G. longipalpis*, and *G. m. submorsitans* display step suitability increases within moderate thermal ranges, followed by declines at higher temperatures, consistent with physiological constraints on tsetse survival. Soil moisture (totSmoist22) often produces positive responses at intermediate to high moisture levels, aligning with the need for humid microhabitats that support adult activity and pupal development.

### Tsetse fly habitat suitability using ensemble modelling approach

#### A. Riverine tsetse flies

*G. tachinoides* showed high habitat suitability in the northwest area and small areas in the south Vallée du Bandama district (Fig. 10). Lagunes district also has small areas of very high habitat suitability on its border with Goh-djiboua. For *G. p. palpalis*, the results showed a high habitat suitability in areas extending from central to the southern regions, in Comoe, Lagunes, Bas-sassandra, western Sassandra-marahoue districts and southern Vallée du Bandama district. The study shows very high habitat suitability for *G. p. gambiensis* in the northern regions, particularly in Savanes district. Whereas the model for *G. pallicera* indicated high suitability in some areas of Bas-Sassandra, and western Sassandra-marahoue.

**Fig. 10.**
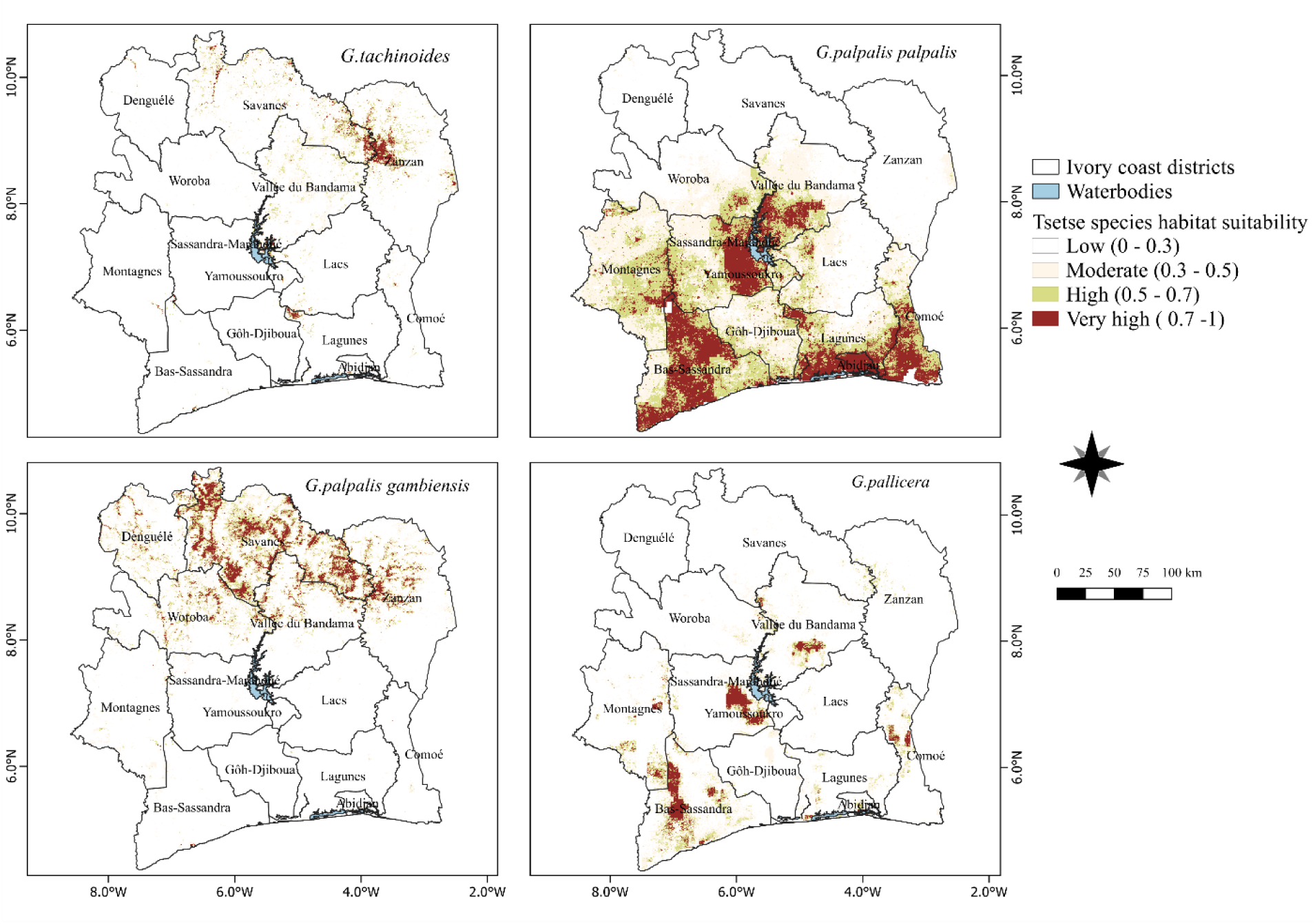
Habitat suitability maps for riverine tsetse fly species (*G. tachinoides*, *G*. *p. palpalis*, *G. p. gambiensis*, and *G. pallicera*) in Côte d’Ivoire generated using a weighted average ensemble species distribution modelling approach.

#### B. Forest flies

The results indicated that *G. medicorum* exhibits high habitat suitability in the Zanzan district, with smaller pockets of suitable habitats occurring in the central part of the Vallée du Bandama district (Fig. 11). Similarly, *G. n. nigrofusca* was predicted to occur in moderately to highly suitable habitats across the southern regions, especially within coastal forest ecosystems and other humid environments.

**Fig. 11.**
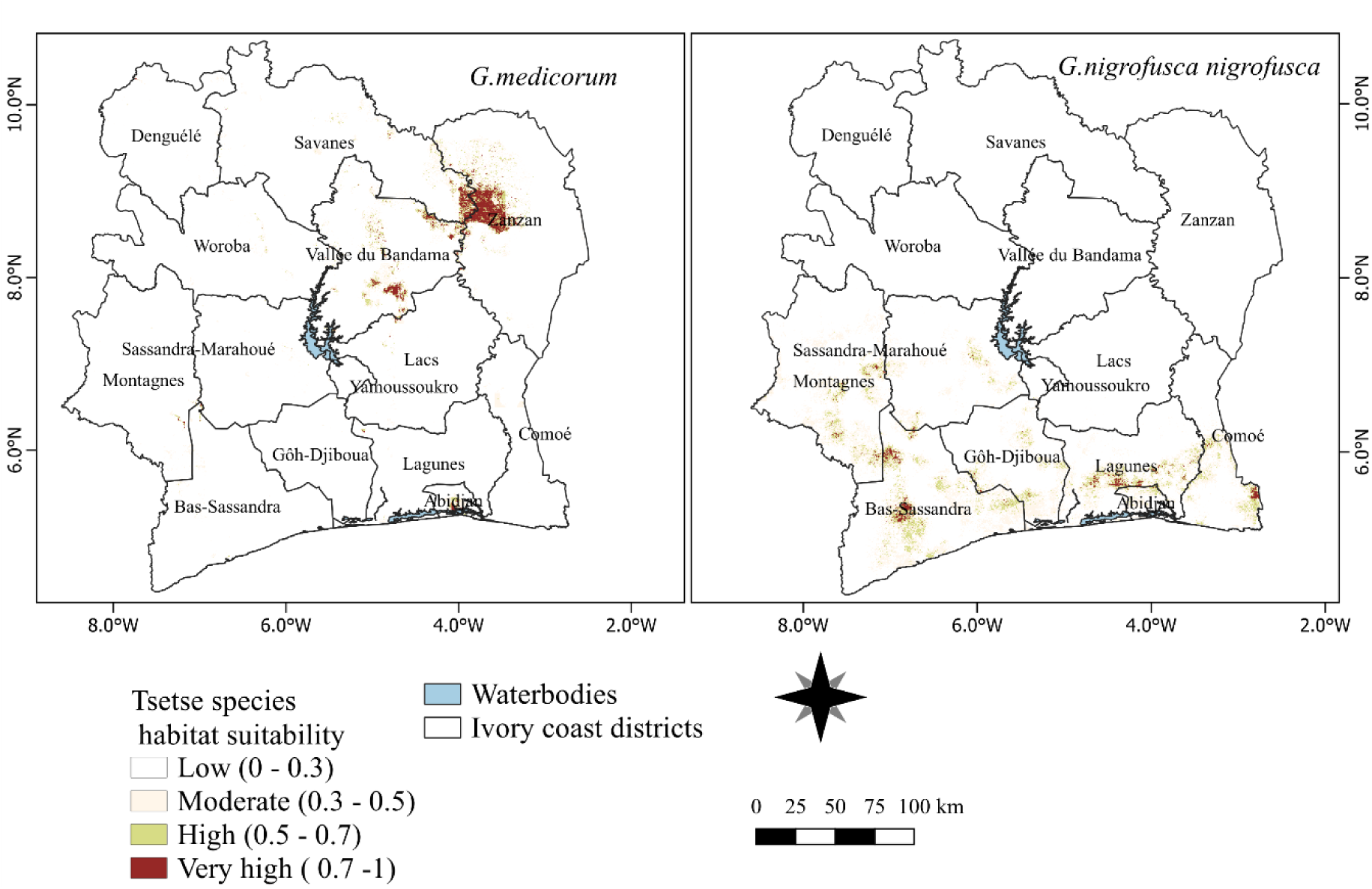
Habitat suitability of forest dwelling tsetse species (G. *medicorum* and G. *n.nigrofusca*) in Côte d’Ivoire generated using a weighted average ensemble species distribution modelling approach.

#### C. Savannah tsetse flies

Patches of suitable habitat for *G. longipalpis* was concentrated in the central-eastern regions of Côte d’Ivoire (Fig. 12), with additional patches of moderate habitat suitability distributed across other parts of the country, particularly in the western Zanzan, and Vallée du Bandama districts. Conversely, *G*. *m. submorsitans* exhibited a more spatially fragmented pattern of habitat suitability, characterized by isolated areas of high to very high suitability, most notably in the eastern Zanzan and north-west Lagunes districts.

**Fig. 12.**
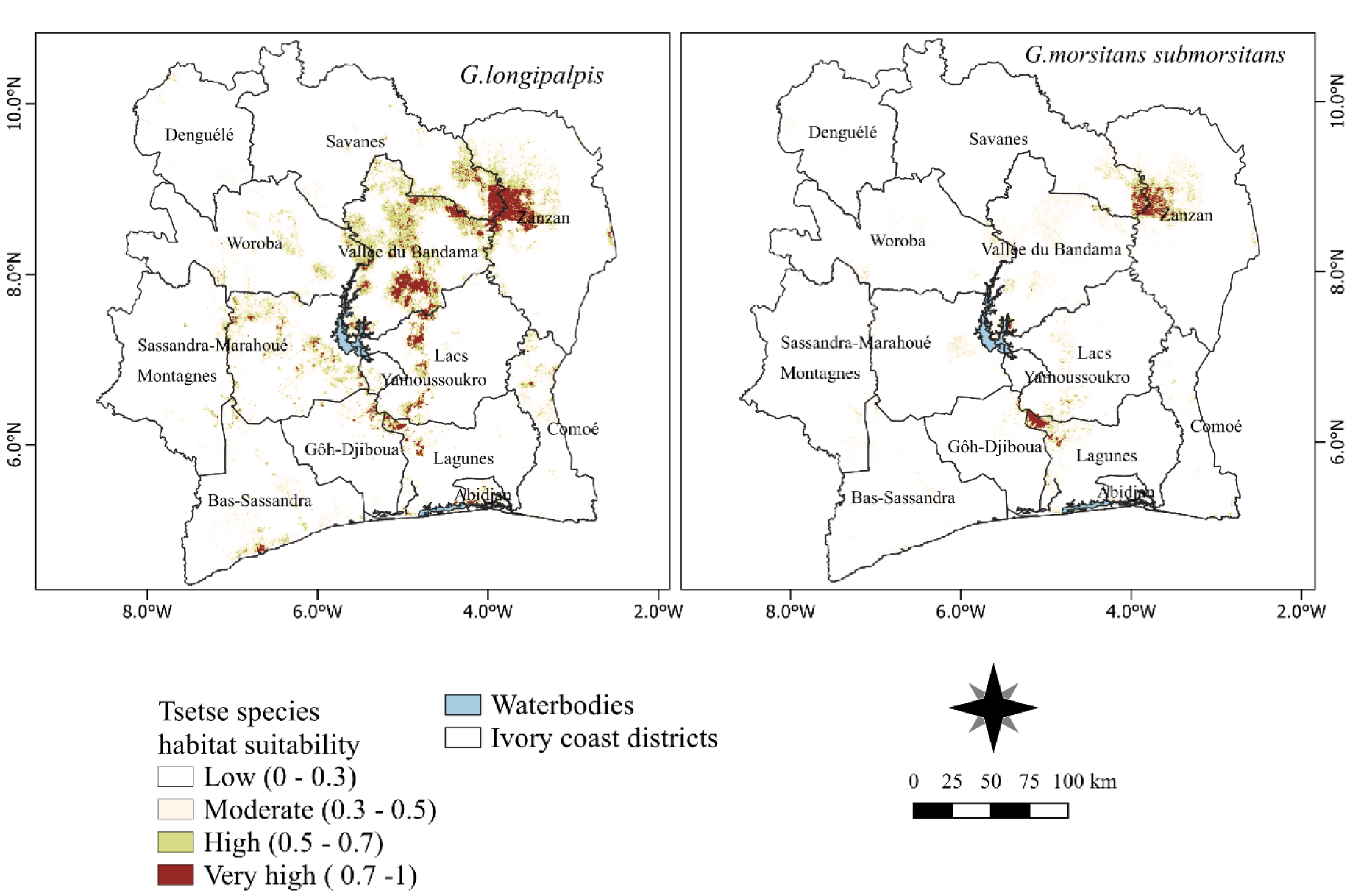
Habitat suitability of two savannah tsetse fly species (*G*. *longipalpis* and *G*. *m. submorsitans*) in Côte d’Ivoire generated using a weighted average ensemble species distribution modelling approach.

#### D. Ensemble species distribution model at genus level (*Glossina spp.*)

These ensemble habitat suitability models were generated from predictions from individual models using TSS-weighted mean scores, excluding underperforming models. At the genus (*Glossina*) level, areas of high habitat suitability were concentrated in the north-eastern, central and southern regions of regions of Côte d’Ivoire (Fig. 13). The weighted mean TSS (0.82) ensembles produced highlight consistent spatial predictions, identifying the same areas as highly suitable. This strong agreement indicates that the nine *Glossina* species share broadly similar ecological niches and overlapping areas of high habitat suitability.

**Fig. 13.**
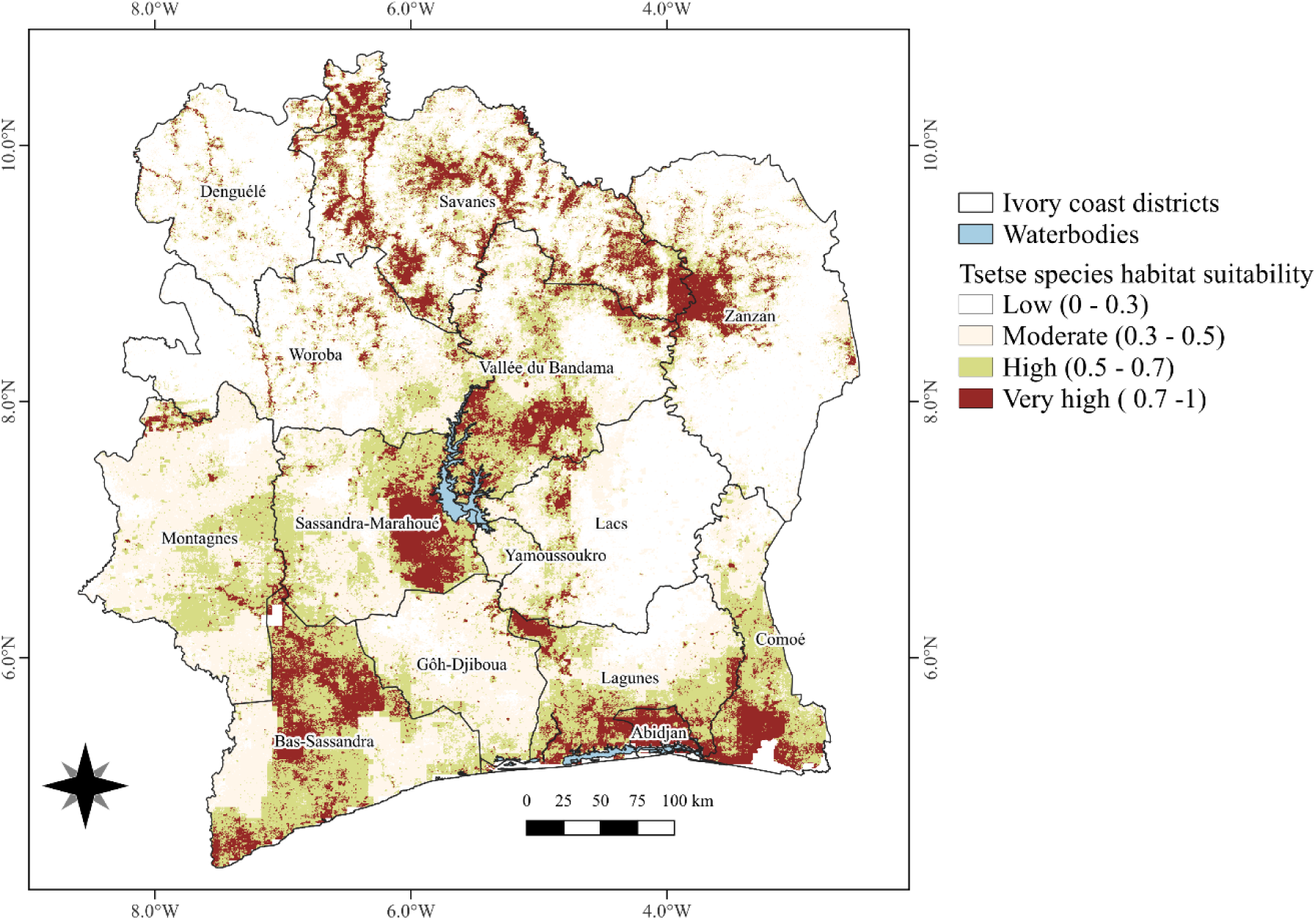
Habitat suitability of tsetse fly (*Glossina* spp.) across Côte d’Ivoire generated using an ensemble machine learning modelling approach based on true skill statistic [TSS]-weighted mean scores.

The predicted habitat suitability was classified into four categories: low (120,212.837 km²), moderate (89,173.312 km²), high (42,067.581 km²), and very high (7,510.996 km²) suitability. Collectively, the high and very high suitability classes encompassed 49,578.577 km^2^, representing the core ecological niches of *Glossina* spp. and, therefore, areas potentially at elevated risk of transmission of both AAT and HAT in Côte d’Ivoire. As shown in Table 4, the largest extents of high and very high suitability were found in the districts of Savanes (5,172.33 km²), Bas-Sassandra (4,813.4km²), Vallée du Bandama (3,041.24 km²), and Sassandra-Marahoué (2,935.77 km²). Lagunes, Zanzan, and Comoé districts also exhibited higher areas of suitable habitat across central and western Côte d’Ivoire, highlighting their importance as priority regions for AAT surveillance and vector control.

**Table 4.** Area (km^2^) of tsetse fly (*Glossina spp*.) habitat suitability across districts of Côte d’Ivoire estimated using an ensemble machine learning modelling approach based on true skill statistic (TSS)-weighted mean scores.

| District | Area (km<sup>2</sup>) |
| --- | --- |
| Abidjan | 834.18 |
| Bas-Sassandra | 4,813.4 |
| Comoé | 1,919.14 |
| Denguélé | 314.26 |
| Gôh-Djiboua | 210.33 |
| Lacs | 555.45 |
| Lagunes | 2,132.07 |
| Montagnes | 597.68 |
| Sassandra-Marahoué | 2,935.77 |
| Savanes | 5,172.33 |
| Vallée du Bandama | 3,041.24 |
| Woroba | 671.67 |
| Yamoussoukro | 38.22 |
| Zanzan | 1,552.07 |

The spatial distribution of suitability varied substantially among Glossina species (S1 appendix, Table S4). Among the individual species, *G. palpalis palpalis* had the largest combined area classified as high and very-high suitability (37,344), followed by *G. palpalis gambiensis* (16,359) and *G. longipalpis* (8,837). *G. palpalis palpalis* also recorded the largest area under the very-high suitability category (3,943), closely followed by *G. palpalis gambiensis* (3,799). In contrast, G. nigrofusca had the smallest extent of very-high suitability (64).

## Discussion

Our results provided an in-depth assessment of the ecological suitability and potential distribution of *Glossina* spp, savannah (*G. longipalpis* and *G. m. submorsitans*), riverine (*G. tachinoides*, *G. p. palpalis*, *G. p. gambiensis*, and *G. pallicera*) and forest species (*G. medicorum* and *G. nigrofusca nigrofusca*) in Côte d’Ivoire thanks to the 2018-2023 occurrence data from the national tsetse atlas (24). The high predictive accuracy of the models, reflected by strong AUC 0.80 and TSS 0.83 scores, demonstrates their reliability in estimating current tsetse habitat suitability across diverse ecological zones of the country. The variability in model performance was not only among the SDM algorithms but also across different *Glossina* s*pp*. emphasizing the importance of algorithm selection in ecological modeling and species distribution studies to support informed decision-making for human and animal health from HAT to AAT.

At the species level, the observed variable-importance patterns largely reinforce genus-wide ecological drivers while also highlighting important interspecific differences related to habitat preference and ecological adaptation. The *Glossina spp.* response curves indicated that minimum LST plays a key role in determining habitat suitability, with temperatures between 280–290° K (7–17 °C) and increasing value from 7°C increases the environmental suitability. Temperatures below this range are linked to reduced suitability, while values exceeding 300 K (∼27 °C) are associated with high suitability, followed by a slight decline at the extreme upper range likely reflecting physiological stress and reduced survival.

Human population density showed a positive relationship with *Glossina spp*. suitability in Côte d’Ivoire, particularly in historical HAT foci such as Bonon, where tsetse abundance (in particular the HAT vector *G. p. palpalis*) is strongly influenced by human activities occurring in landscapes where riverine vegetation overlaps with human–vector contact zones (71). This pattern is consistent with the strong influence of LULC classes associated with human settlements and infrastructure. However, previous studies (72) indicated that very high human population densities may reduce suitability due to habitat degradation and wildlife disappearance, particularly for species of the fusca group (73) and the *morsitans* group, whereas the opposite trend has been observed for *G. p. palpalis* (74). In fact, there is only a limited increase and very quickly a plateau for fusca group flies with the human population (with even a decrease and then increase before plateauing for *G. medicorum*), a decrease for *G. morsitans. submorsitans*, and only really a large increase for *G. p. palpalis* and a small decrease for *G. p. gambiensis*. Hence, the lack of any strong positive correlation and even sometimes somewhat negative for most species and the strong positive correlation for *G. p. palpalis* is associated with the larger sampling effort for *G. p. palpalis* linked to HAT may drive our results to some extent. Indeed, tsetse traps are often deployed in close proximity to human settlements, particularly in areas targeted for HAT surveillance. Consequently, tsetse occurrence records are inherently associated with human presence, introducing a sampling bias that may not fully capture the species’ ecological niche. As a result, the observed distribution may represent only a subset of the realized niche, potentially overlooking suitable habitats in areas that have not been systematically surveyed.

Across multiple species, vegetation greenness, captured through NDVI-derived metrics, emerges as a key predictor, particularly for *G. longipalpis*, and *G. n. nigrofusca*, which were exclusively captured in protected areas (24). High NDVI values are typically associated with favorable microclimatic conditions (20,46,75), including reduced temperature extremes and increased humidity, which are essential for adult survival and pupal development (15,76).

The major LULC classes contributing to tsetse habitats include a mix of natural forests, agroforestry systems, and open landscapes. Riverine and forest-associated tsetse fly species presence are primarily linked to open forests, gallery forests, secondary or degraded forests, mangroves, swamp forests, and forest plantations, which provide essential shade, humidity, and breeding sites (16,77). Simultaneously, agroforestry plantations such as coffee, cocoa, and rubber offer positive suitable microhabitats for riverine tsetse (*G.palpalis sensu lato*). However, new farming practices, characterized by the excessive use of pesticides, have made these areas hostile to tsetse fly (78), agricultural lands, fallow fields, wooded savannas, shrublands, and grasslands have a positive influence on species’ occurrence depending on vegetation cover, proximity to water bodies, and livestock presence.

The response curves revealed nonlinear thermal relationships, with species such as *G. medicorum*, *G. longipalpis*, and *G. m. submorsitans* exhibiting peak habitat suitability within intermediate temperature ranges, followed by a decline at higher temperatures. This pattern is consistent with established evidence that tsetse flies function within relatively narrow thermal limits, beyond which survival, fecundity, and pupal development are adversely affected (49,79).

Moderate human population densities likely maintain fragmented habitats suitable for tsetse reproduction and dispersal by preserving vegetation patches and mixed farming systems that facilitate tsetse persistence. Recent studies shown that tsetse are commonly associated with wildlife-livestock-human interface areas, where interactions among wildlife reservoirs, domestic livestock, and human populations facilitate the persistence and transmission of trypanosomiasis (80,81). In Côte d’Ivoire, previous research has documented a strong association between pig keeping and *G. palpalis palpalis* occurrences (82–84). In this study, we interestingly observed that human population density positively influences *G. p. palpalis* suitability, likely reflecting the persistence of mixed-use landscapes that provide resting habitats and host accessibility, which may have been conducive to an increase in HAT infection. In addition, high pig densities have a negative correlation with *G. p. palpalis* suitability, possibly due to more intensive livestock management practices such as restricted free-ranging or insecticide treatment [protective nets], which could limit tsetse access to hosts. For other species, including*. G. longipalpis*, *G. morsitans submorsitans*, *G. palpalis gambiensis*, and *G. tachinoides*, we observed a positive correlation with pig density, suggesting that livestock presence provides accessible blood-meal sources that increase habitat suitability.

In northern Côte d’Ivoire, cattle production plays a central role in rural livelihoods and agricultural intensification, particularly within the cotton basin where AAT remains a major constraint to livestock rearing and cotton production. This region is predominantly affected by trypanosome species such as *Trypanosoma vivax*, *T. congolense*, and *T. brucei*, transmitted mainly by savannah and riverine tsetse species including *Glossina morsitans submorsitans*, *G. palpalis gambiensis*, and *G. tachinoides* (85). The negative association between *Glossina spp*. suitability and high livestock densities likely reflects the fact that, in Côte d’Ivoire, high-intensity livestock production systems are often linked to habitat degradation and to targeted vector-control interventions, both of which can reduce tsetse fly populations. In contrast, the relationship between cattle density and *Glossina spp.* suitability appears non-linear, with low-to-moderate cattle densities (0–40 head/km2) associated with increased suitability. At the species level, suitability increased with cattle density for *G. longipalpis, G. palpalis gambiensis, G. morsitans submorsitans*, and *G. tachinoides*, whereas suitability decreased with increasing cattle density for *G. fusca fusca, G. medicorum, G. pallicera*, and *G. palpalis palpalis*. These contrasting responses likely reflect differences in ecological preferences and host use, with cattle acting as key blood-meal hosts that can sustain tsetse populations in mixed agro-pastoral systems. Notably, cattle density remained positively associated with suitability in parts of northern Côte d’Ivoire, despite concurrent pressures such as riparian vegetation degradation and intensified AAT control strategies, including insecticide-treated cattle and targeted vector-control campaigns.

The small ruminants, sheep and goats, have been identified as key in the long-term field studies in northern Côte d’Ivoire. It has been shown that sheep are exposed to trypanosome infections alongside cattle under natural grazing systems (32), confirming their participation in the transmission cycle, consistent with the strong influence of tsetse fly suitability. Additionally, multi-host epidemiological surveys in endemic foci such as Bonon and Sinfra have detected trypanosome infections in sheep, although the infection rates were higher in cattle and pigs, supporting their role as potential reservoirs in AAT-endemic landscapes (32).

Distances to protected areas and water bodies have a strong negative relationship with tsetse fly suitability. These environments provide favorable humidity, temperature regulations, and suitable breeding substrates critical for adult survival and larval development (15,86). Distance to water bodies emerges as a key predictor for riverine species such as *G. palpalis palpalis* and *G. tachinoides*, as well as for the forest-associated *G. nigrofusca nigrofusca*. This aligns with their ecological dependence on humid environments, where proximity to rivers, wetlands, and shaded riparian vegetation provides suitable microhabitats (73). In West Africa, riverine systems often act as corridors for tsetse dispersal, further reinforcing this relationship (87,88).

As indicated by the variable importance results, climatic factors such as LST, vegetation, LULC cover in Côte d’Ivoire are critical determinants shaping tsetse habitat suitability. Therefore, incorporating mechanistic and mathematical models alongside SDMs or within hybrid SDMs could deepen our understanding of how ongoing habitat fragmentation and landscape transformations in relation to climate change affect tsetse fly ecology, AAT transmission dynamics and risk for HAT transmission rebounds. Recent studies reinforce this approach, e.g., Helikumi and Mushayabasa (89) applied a compartmental modelling framework to human-cattle-wildlife-tsetse systems, demonstrating how host composition and land-use patterns influence transmission dynamics; (90) employed a mechanistic population-dynamics model of *G. pallidipes* in Zimbabwe, linking environmental changes to variations in vector abundance; (91) demonstrated that incorporating species phenology, life-history traits, and occurrence records into hybrid frameworks captures both the ecological processes driving distribution and the statistical relationships with environmental predictors, and (92) integrated machine learning and genetic connectivity analyses to improve spatially explicit vector control planning.

It is also useful to note that our model outputs are influenced by the availability and quality of tsetse occurrence data and environmental covariates to predict suitable areas. Indeed, the current occurrence dataset for Côte d’Ivoire is somewhat biased toward riverine tsetse species (*G. palpalis palpalis* and *G. palpalis gambiensis*), reflecting a historical focus on entomological data gathering aiming at HAT surveillance (24). Species associated with AAT, such as *G. morsitans submorsitans* and some of the forest flies, such as *G. fusca fusca* are underrepresented, and further surveillance efforts will improve our prediction of the spatial risk for AAT, especially in the cotton basin in the North. Besides, habitat fragmentation may alter both vectors and host populations, thereby influencing disease dynamics in ways not easily represented in SDMs (93).

Finally, Côte d’Ivoire achieved elimination of HAT as a public health problem through sustained medical surveillance, diagnosis and treatment, complemented by targeted vector control. Subsequent modelling of surveillance data provided additional evidence of declining transmission and supported assessment of progress towards interruption of transmission (4). Building on this success, informed and strategic decision-making supported by the species distribution models presented here, together with future epidemiological and livestock production studies, should thus be encouraged to achieve sustainable control of AAT. The habitat suitability maps generated in this study provide a valuable evidence base for identifying priority areas for targeted entomological surveillance, particularly in regions characterized by high predicted suitability and substantial livestock populations. Integrating tsetse maps into decision-support frameworks can strengthen the ecological basis for HAT and AAT control basis for implementing the Progressive Control Pathway (PCP) for both HAT and AAT. Furthermore, combining these maps with socio-economic indicators such as livestock ownership, poverty levels, and access to veterinary healthcare - would facilitate a ‘One Health’ approach enabling control programs to prioritize interventions in the most vulnerable communities and maximize the efficiency and sustainability of disease control (23,94).

## Conclusions

The ecological suitability and potential distribution of eight *Glossina* species across Côte d’Ivoire were predicted, identifying the key environmental and anthropogenic drivers shaping their habitats. The high predictive accuracy of the models demonstrates their reliability, while response curves reveal that tsetse fly presence is influenced by optimal LST, human and livestock density, proximity to water bodies, and LULC patterns. Our findings provide an evidence base to support policymakers and decision-makers in identifying priority areas for targeted surveillance, vector control, and, where feasible, the local elimination of tsetse populations, therefore contributing to the sustained interruption of HAT transmission and improved control of AAT.

## Supporting information

- S1 Appendix_Variable importance results
- S2 Appendix _Variable response curves
- S3 Appendix_Species distribution modeling code
- S1 Dataset_Species occurrence data
- S1 Folder_Environmental covariates data

## Data availability

All relevant data supporting the findings of this study are fully available within the manuscript and its Supporting Information files

## Acknowledgments

The authors gratefully acknowledge the financial support of the European Union’s Horizon 2020 research and innovation programme under grant agreement n°101000467, acronym ‘’COMBAT’’ (Controlling and Progressively Minimizing the Burden of Animal Trypanosomosis).

FAO support was provided within the framework of the Programme Against African Trypanosomosis (PAAT), and it was enabled by FAO participation in the project COMBAT. Furthermore, FAO assistance was also supported by the project ‘Disease intelligence & modelling for progressive control of animal trypanosomosis in Africa’ (DIMCAT), funded by the Gates Foundation (INV-067581). The authors sincerely appreciate the financial support provided by the Swedish International Development Cooperation Agency (Sida); the Swiss Agency for Development and Cooperation (SDC); the Australian Centre for International Agricultural Research (ACIAR); the Government of Norway; the German Federal Ministry for Economic Cooperation and Development (BMZ); and the Government of the Republic of Kenya.

